# MolecularWebXR: Multiuser discussions about chemistry and biology in immersive and inclusive VR

**DOI:** 10.1101/2023.11.01.564623

**Authors:** Fabio J. Cortés Rodríguez, Gianfranco Frattini, Sittha Phloi-Montri, Fernando Teixeira Pinto Meireles, Danaé A. Terrien, Sergio Cruz-León, Matteo Dal Peraro, Eva Schier, Kresten Lindorff-Larsen, Taweetham Limpanuparb, Diego M. Moreno, Luciano A. Abriata

**Author notes:** Corresponding autor.

## Abstract

MolecularWebXR is a new website for education, science communication and scientific peer discussion in chemistry and biology, based on modern web-based Virtual Reality (VR) and Augmented Reality (AR). With no installs as it is all web-served, MolecularWebXR enables multiple users to simultaneously explore, communicate and discuss concepts about chemistry and biology in immersive 3D environments, by manipulating and passing around objects with their bare hands and pointing at different elements with natural hand gestures. User may either be present in the same real space or distributed around the world, in the latter case talking naturally with each other thanks to built-in audio features. Although MolecularWebXR is most immersive when running in the web browsers of high-end AR/VR headsets, its WebXR core also allows participation by users with consumer devices such as smartphones, possibly inserted into cardboard goggles for deeper immersivity, or even in computers and tablets. MolecularWebXR comes with preset VR rooms that cover topics from general, inorganic and organic chemistry, biophysics and structural biology, and general biology; besides, new content can be added at will through moleculARweb’s PDB2AR tool or by contacting the lead authors. We verified MolecularWebXR’s ease of use and versatility by people aged 12-80 years old in entirely virtual sessions or in mixed real-virtual sessions at various science outreach events, in courses at the bachelor, masters and early doctoral levels, in scientific collaborations, and in conference lectures. MolecularWebXR is available for free use without registration at https://molecularwebxr.org, and a blog post version of this preprint with embedded videos is available at https://go.epfl.ch/molecularwebxr-blog-post.

## Introduction

In education and also in the daily work in the chemical and biological sciences, the human ability to visually grasp and communicate the details of objects that are inherently three-dimensional is essential, yet most technical means of presenting and manipulating 3D information are intrinsically two-dimensional.^1^ In fact, today most molecular visualization and manipulation takes place in the form of two-dimensional interfaces and representations such as pictures, diagrams and flat-screen computer graphics. Even the most advanced molecular graphics programs at their core typically display molecules on flat 2D screens, and allow users to interact with the molecular system only through mouse moves and key strokes which are inherently one-handed and very limited in terms of natural interactivity, let alone in allowing concurrent action of multiple users. Attempting to alleviate these drawbacks, over various “waves of hype” on virtual reality (VR) most classical programs for molecular modeling and graphics introduced VR extensions that provide immersive visualization and spatial control on manual operations.^2–4^ In addition, various programs have been developed specifically for manipulating molecules in VR, often with limited functionality compared to the VR versions of more complete molecular graphics software but with the benefit of being much simpler to deploy and utilize. Besides, tests have been conducted to explore the benefits and drawbacks of the technology in real-world utilization, including tests on how multiple users can work concurrently in shared VR sessions.^5^ However, the adoption of VR software for molecular graphics has been very limited, and one could argue that the lack of success is most likely due to a technology that was not ripe enough until this decade.

Since around year 2020, a new wave of VR and also augmented reality (AR) technologies has opened up new frontiers for immersive and interactive learning experiences, with these technologies evolving faster than ever among consumers.^6^ This renewed interest in VR and AR, along with the substantial investment fueled by the idea of building the pieces of a “Metaverse”, has fostered the development of new AR and VR devices that are smaller and more wearable, have sensors that track the environment and the user’s pose with high accuracy, and are fully built into the headset eliminating the need to connect to external computers. Recent AR and VR headsets include built-in Wi-Fi connectivity and web browsing capabilities and are more affordable. For example, immersive devices like Meta’s Oculus Quest 2 and 3, used in this work, are available for USD 300-600 as of June 2024.

Together with the advent of new AR and VR hardware, new programming standards have also evolved coordinated by major vendors, and new software tools have emerged in the last years that allow users to teach, self-learn and work in immersive environments.^7^ In the specific context of chemistry and biology,^8^ stand-alone programs for AR/VR and modules that extend the capabilities of regular computer graphics programs into AR-VR have emerged.^9–14^ Importantly, the most advanced of these AR/VR programs are starting to support at least two-user sessions,^5^ thus slowly starting to enable virtual human interactions and collaborations as required for teaching, collaborative work and discussions, etc.

However, despite the recent emergence of several new programs for immersive molecular visualization and modeling in VR/AR, their adoption still seems very limited. We argue that the main limitation is not only the cost of the AR and/or VR headsets, which has lowered but is still substantial, but also involves the complex nature of the setups involved and the limited cross-device compatibility of the existing programs. This is why over the last 5 years we have advocated for and built purely web-based solutions for molecular graphics,^15^ meant to work “out of the box” without installation, recently capitalizing on the WebXR standard and API.^16^ WebXR-based solutions bring two advantages of unparalleled relevance to adoption and democratization of access to immersive content: (i) high portability across devices and operating systems, from laptops and smartphones to high-end AR/VR headsets; and (ii) low barriers to deployment and availability as all content runs inside web browsers hence requires no installs or updates and is instantly available upon internet connection.^16–18^ Plus, by not requiring any login or registration, the website is used under full privacy (and no files loaded by the users are stored in our server).

With this philosophy in mind, 3 years ago we released moleculARweb, a free platform that offers several activities and playgrounds for chemistry and structural biology education in commodity AR, that is AR that runs on non-specialized devices such as smartphones, computers, and laptops.^19–21^ Next, building on the WebXR standard we extended moleculARweb with PDB2AR, a web tool that allows users to create web-based AR and VR sessions for any kind of consumer device, including basic forms of WebXR-based display. In PDB2AR the virtual objects are generated with VMD,^22^ and as such it supports all its “representations” from simple ball-and-stick models and cartoons to isosurfaces useful to represent electronic orbitals or cryo-electron maps.^23^

Here we introduce MolecularWebXR (Figure 1A), a new platform for education, science communication and scientific peer discussion in topics of chemistry and (mainly structural) biology exploiting immersive VR and AR as supported by the WebXR standard. Being web-served and hence not requiring installation or any additional software, MolecularWebXR enables multiple users to simultaneously explore, communicate and discuss concepts about chemistry and biology in immersive 3D environments by manipulating and passing around objects with their bare hands and pointing at their features with natural hand gestures. Users can be present in the same physical space or distributed around the world, in the latter case talking naturally with each other through the device thanks to built-in audio features. MolecularWebXR offers rooms with pre-built material about a series of topics for chemistry and (structural) biology education, and an empty room that users can populate with custom-made objects from moleculARweb’s PDB2AR app to create personalized VR sessions for classes, demonstrations, and scientific discussions.

**Figure 1.**
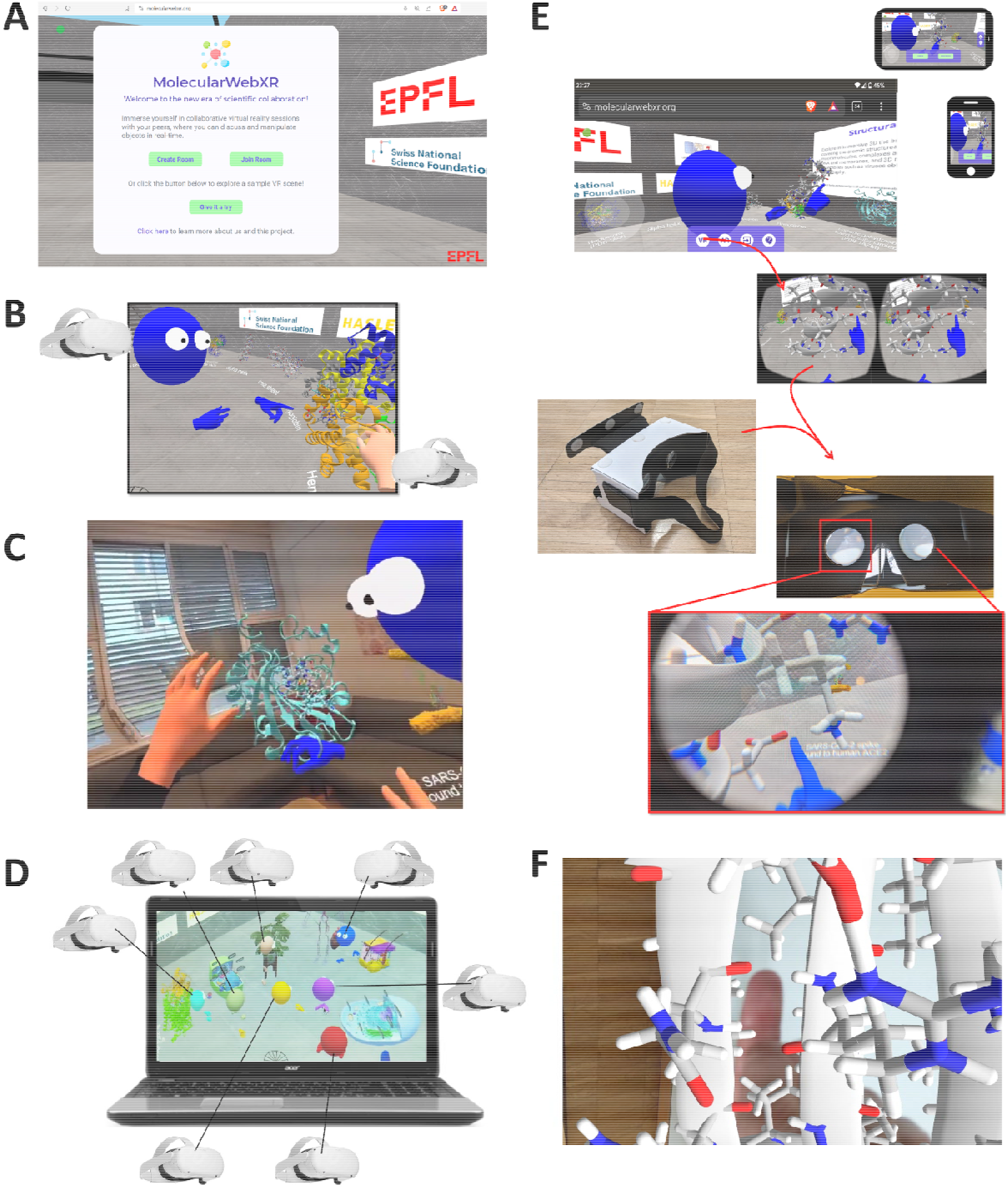
MolecularWebXR as seen in web browsers on various WebXR-capable devices. (A) Welcome page as seen in a laptop. (B) In VR headsets, the 6 degrees of freedom allow users to explore the VR scene simply by walking around and moving their heads and limbs naturally. In addition, users can translate, rotate and scale objects via natural manual operations with their hands or by using the device’s handheld controls. The hands/controls and the heads of participants using VR headsets are broadcast to all other users as simplified avatars. If audio is on, these users can naturally talk to each other and with guests who follow the session from any device can listen to the users in VR. (C) A two-user session seen from the view cast by a user wearing an Oculus Quest 3 (orange hands) while another user (blue avatar) follows closely. (D) A session featuring seven speakers wearing VR headsets as seen from a laptop (looks similar on a tablet). Users can move around the scenes by using the W, A, S and D or the arrow keys if the keyboard (in computers) or a virtual joystick (in smartphones and tablets), and they can direct their looks by using the mouse (computers) or *via* touch gestures (tablets). (E-F) A session with a user in VR mode accessing through a headset (blue avatar) as seen by a user accessing the session with a smartphone. Outside of VR (top) users accessing via smartphones can move around the scene with a virtual joystick and they can gaze *via* touch gestures. In smartphones supporting WebXR (going down through the panel), users can enter WebXR mode and either insert the phone into cardboard goggles to explore the scene via 3 degrees of freedom at the entry point in VR (panel E, without the ability to move around the session but with the option to move their heads to look around naturally) or simply see through the phone for AR (panel F, with six degrees of freedom). For more on how to use MolecularWebXR, see Figures S1-S3.

Inside MolecularWebXR sessions, users wearing VR headsets are displayed with their heads and hands reflecting their natural poses and moves; they can grab objects to move them and resize them in space, point with their hands in a natural fashion, and communicate with each other by talking naturally directly through the headset’s microphone, in VR (Figure 1B) or AR (Figure 1C). Users without access to hardware specialized for VR can still follow the sessions from their laptops, tablets or smartphones (Figure 1D-F). Moreover, the latter supports immersive VR modes with 3 degrees of freedom (rotations) by using cardboard-made goggles (Figure 1E), and through-screen AR with 6 degrees of freedom (Figure 1F).

Throughout the article we showcase example applications of MolecularWebXR as either entirely virtual sessions or mixed real-virtual sessions, in VR and in AR, deployed in science outreach days at our institutions, student instruction at courses of varied levels, scientific collaboration, and at conference lectures.

### System

MolecularWebXR relies on WebXR, an API and specification for web content and web apps to interface with mixed reality hardware and supported by all major vendors, thus easily enabling cross-device compatibility.^16^ This API automatically parses the device’s input capabilities into standardized events and mechanisms that are fed into the web browser, for which the software is written in JavaScript at the core. A server, running in Node.js, centralizes the creation and management of rooms where VR sessions take place. The exact VR devices used in the experiences presented throughout the figures of this article were the Oculus Quest 2, Meta Quest 3 and Meta Quest Pro, all with the hand tracking feature enabled. We also allow object manipulation with handheld controls in these 3 devices, and we verified proper working of the website in the App Vision Pro, Oculus Quest 1 and HTC Vive Pro, the latter two only handheld controls.

By design, access to MolecularWebXR is highly democratized, as the web standard ensures that the software works out of the box in the web browsers of all kinds of devices from high-end VR headsets to smartphones, tablets and computers, leaving no one out as discussed above and exemplified throughout all figures of this paper. In high-end VR headsets, users can grab objects and pass them around with their hands or controls, zoom them in or out with natural gestures, use their hands to point at objects, and freely move around scenes, in AR or VR (Figure 1B-D). Users accessing with headsets are displayed as hands-and-head avatars that all other users can see. In modern smartphones, users can move around the scene with a joystick located on the bottom left and choose where to gaze by touching on the screen (Figure 1E top, where the joystick is the grey circle on the bottom left). They can also access sessions in immersive VR by using cardboard goggles, without the ability to grab objects as smartphones do not (yet) offer built-in hand tracking but with full capability to see the scene and the other users as well as hearing the conversations and talking. A limitation of smartphones in the goggle-assisted VR mode is that they only offer 3 degrees of freedom, allowing 360º visualization but not displacements around the scene, which can only be achieved outside of VR mode by using the joystick. However, modern smartphones also allow see-through AR with 6 degrees of freedom as in Figure 1F. Last, in tablets, laptops and other kinds of computers there is naturally no immersive AR or VR of any type; however, users can move around the VR scenes by using the arrow (or W, A, S and D) keys and mouse or touch gestures and they can see and hear all users who are inside AR or VR.

### Using MolecularWebXR

To access MolecularWebXR, users must direct the web browsers of their devices to https://molecularwebxr.org/. On first entry with a given device, users must allow audio functions to support talking over the internet, if this feature is to be used. No login or registration is required. For details on how to use MolecularWebXR, see Figures S1-S3.

Once in the main hall (Figure 1A), the user can create a new room or join an existing one by using a unique code provided by the person who created it, whom we refer to as the *Admin*. When a user creates a new room, s/he becomes its *Admin* and obtains codes to invite *VR-active* and other users who can act as guests, *i*.*e*. who can follow the presentation but only passively. *VR-active* users can enter VR if they use a WebXR-capable device and can grab virtual objects with their hands or controls if they are using a VR headset and if the *Admin* has enabled object grabbing. *VR-active* users can also talk to each other and to the *Admin*. All users can move around the VR scene and listen to the *Admin* and *VR-active* users, but those in passive mode cannot grab objects or talk. This distinction between different classes of users allows sessions to be set up in different formats; for example, as discussions where *VR-active* users can present and discuss while other users follow passively; or as courses/lectures/presentations where a single *Admin* or *VR-active* user presents to various passive users.

Importantly, VR sessions running in MolecularWebXR can accommodate users located in the same physical space or accessing from remote locations (Figure 2A and 2B). In the latter case, the audio features are essential to allow for natural conversation and discussion. If all users are located in the same physical space, the audio features should rather be turned off. For mixed locations, audio features can be kept at the *Admin* level, but ensuring that individuals located in the same physical space turn off all but one of their personal audio inputs.

**Figure 2.**
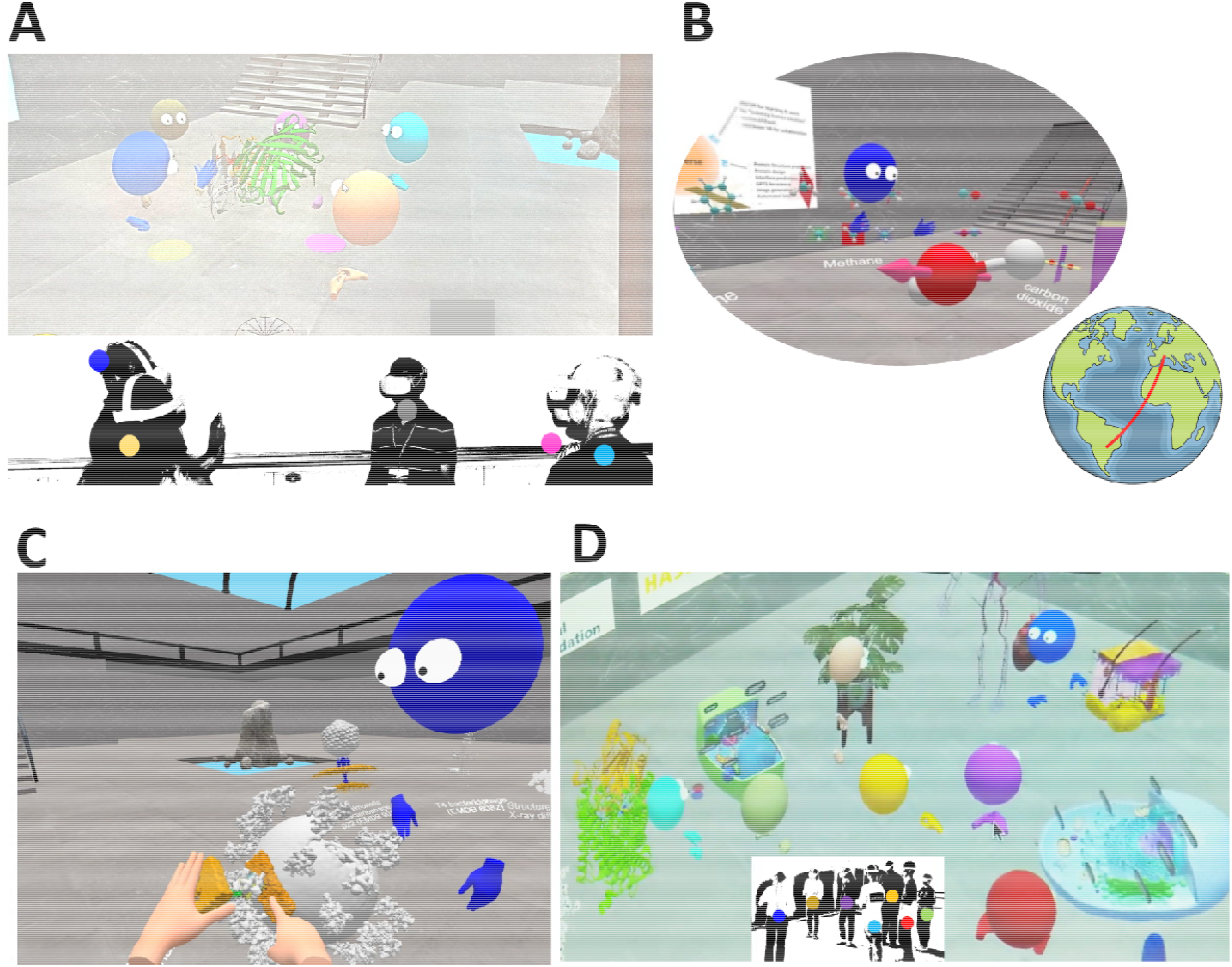
Real vs. virtual presence in MolecularWebXR, and example content. (A) Five users attending a VR session where the avatar in blue is describing the structure of a protein complex. Example taken from an application of the website to a subset of talks during a structural biology conference. The safety spaces of the 5 VR headsets were synchronized to optimize match between real and virtual worlds, and audio features were off. Other attendees of the conference could follow the talks by projecting the view of a sixth user accessing through a laptop. (B) A teacher in Rosario, Argentina (blue avatar) teaching his students from inside a room populated with VR content about the symmetry elements of molecules, as seen by a visitor accessing in a VR headset from Lausanne, Switzerland, over 11,000 km away. Students in this case followed the teacher’s presentation through the view of a third user who accessed the session with a laptop and projected its view on a widescreen. (C) Two users inside the same VR session, as seen from the viewpoint of the orange avatar as she/he is aligning a model of the SARS-CoV2 Spike protein bound to an ACE2 receptor to the electron density of a Spike protein protruding from the viral particle reported from cryo-electron tomography and subtomogram averaging deposited as EMDB 30430^24^. (D) A session deployed over an open science day at EPFL using MolecularWebXR with seven people in the VR room, on content prepared by combining custom representations of molecules created by PDB2AR with models obtained from Sketchfab.com in free or paid forms (this room is not available on the website as it contains purchased objects). For photograph in C and D colors on the users were removed to mask their identities and we have overlaid circles whose colors match the corresponding avatars as seen in the projected views.

When a session consists of users in the same physical space, it is important that the safe-space of all VR devices (the “guardians” in the jargon of Oculus/Meta products) is set up in the same way, so that the relative positions of different users match in the real and virtual spaces. With the right setup, users can feel their physical proximity and talk directly to each other in a very natural way, in the same real space but handling the virtual objects and not seeing their real bodies but their avatars.

We note that bandwidth consumption is high when the VR objects are downloaded upon the user entering the room. Once the room is ready, very high-speed Wi-Fi is not needed but it is important to have a stable internet connection to keep updates fluid as the different users move objects and themselves around the virtual room. We note that no video is transmitted between users but just the quaternions that describe object positions, orientations and scales of the VR objects and of the hands and head avatars of all users inside VR.

We have had up to 8 simultaneous *VR-active* users inside VR, an *Admin* running on a laptop and two passive users following the sessions online, without apparent lag (all 8 VR devices and the *Admin* were on the same Wi-Fi network).

### Content and example applications

When the *Admin* user creates a session, she/he can choose to use a preset room containing content that we have prepared specifically to cover certain topics of chemistry and structural biology expected to benefit largely from deep 3D visualization, or from an empty room where objects can be added at will (Figure 3). Fully customized content can be created via PDB2AR (https://molecularweb.epfl.ch/pages/pdb2ar.html) (Figure 3A). The preset content covers topics on molecular structure (Figure 3B-F), atomic and molecular electronic orbitals (Figure 3C), symmetry elements in molecules of the main point groups (Figure 3B), crystal latices and atomic arrangements in simple materials (Figure 3D), introductory structural biology (Figure 3F), and 3D views of cellular compartments and viruses obtained by cryo-electron tomography (Figure 3G-I).

**Figure 3.**
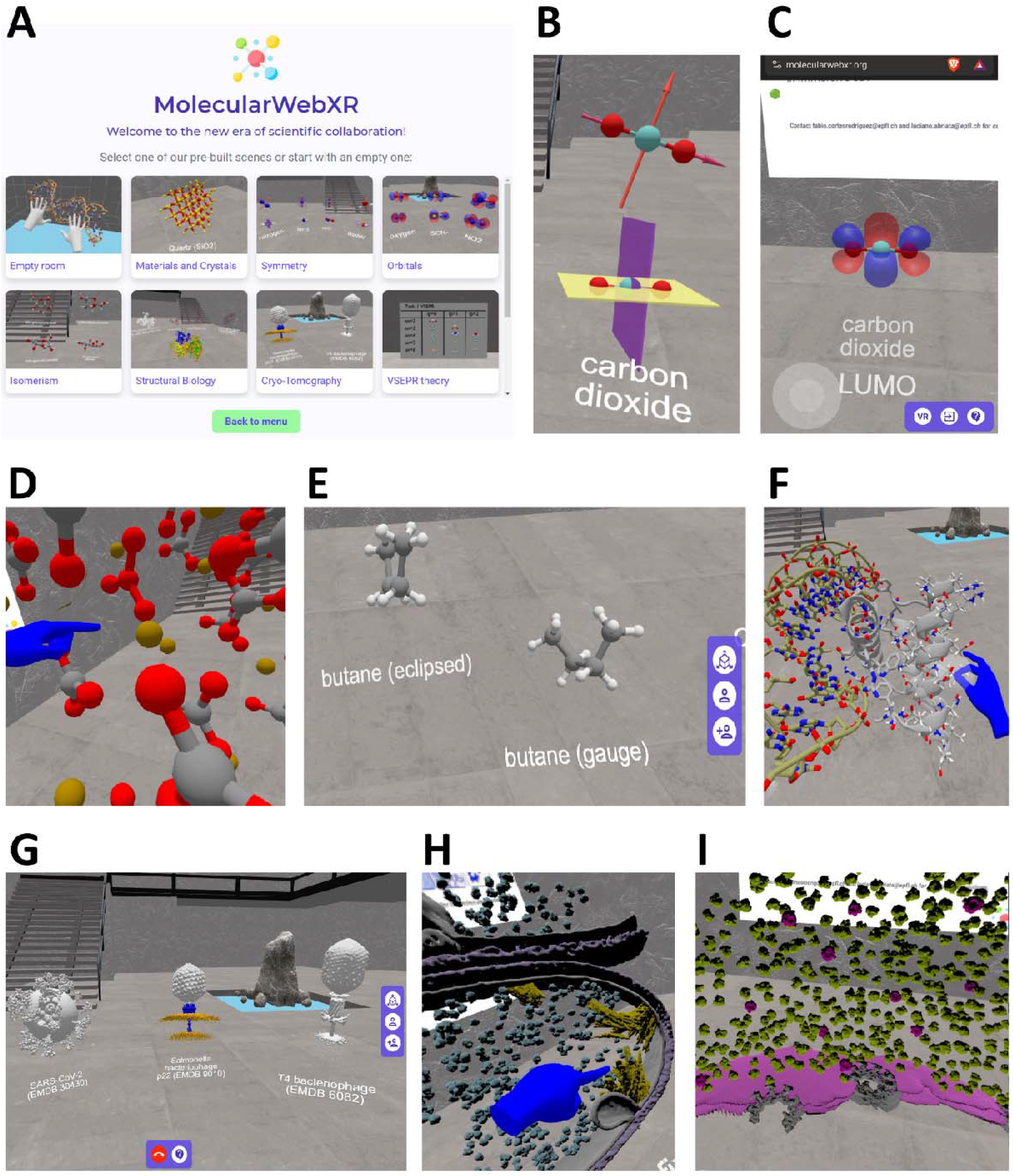
Examples of 3D content running in VR sessions inside MolecularWebXR. (A) All the rooms and pre-built content available at MolecularWebXR at launch. Besides the empty room to be populated with objects from PDB2AR, we offer rooms displaying the symmetry elements of molecules from the main point groups, the frontier molecular orbitals most widely studied at university level, examples on isomerism, example structures of materials and crystalline arrangements, example structures of biological macromolecules, and examples of subcellular structures imaged in 3D through cryo-electron tomography. (B) Small cut from a laptop screen around CO_2_’s symmetry elements, from the room on molecular geometry. (C) Screenshot from a smartphone in portrait orientation showing O_2_’s LUMO, from the room on molecular orbitals. (D) Zoom inside VR on the local geometry around Ca^2+^ ions in calcite, CaCO_3_, with the viewer’s index finger pointing at a Ca^2+^ ion. (E) Models of butane in eclipsed and gauge conformations, from the room on isomerism. (F) Superimposing by hand an atomically detailed model of an alpha helix onto a helix in a cartoon-only representation of a transcription factor bound to DNA. (G) A SARS-CoV-2 particle, a *Salmonella* bacteriophage punching through the two bacterial membranes (orange) with its needle (blue), and a T4 bacteriophage, all retrieved from the indicated the EMDB entries displayed in the labels, as seen in the room about cryo-tomography for biology. (H) Also from the room on cryo-tomography for biology, cut of *Thermoanaerobacter kivui* cell showing the carbon-fixing organelles in yellow, the membrane and some of its invaginations in grey, and the S-layer in purple (Model from ref. ^25^). (I) Another object in the room on cryo-tomography for biology: 3D landscape around the nuclear envelope for a *S. pombe* cell. The model was obtained from ref. ^26^ as detailed in ref.^27^ and shows the nuclear envelope (purple) with some nuclear pore complex subunits (grey), 80S ribosomes (green) and fatty acid synthases (magenta).

Immersive 3D visualizations are particularly well suited for analyzing *in situ* cryo-electron tomography data to unravel the spatial interactions that underlie cellular organization. In fact, researchers working at the frontier of automatic structure identification have shown great enthusiasm for the possibilities offered by MolecularWebXR. Indeed, some groups have contributed with cellular 3D reconstructions, and we are open to add more material on request. Figure 3H shows a map of *T. kivui* cells. The model depicts the plasma membrane, S-layer, and the unusual filamentous enzyme that drives CO_2_ fixation.^25^ Figure 3I shows a 3D landscape around the nuclear envelope of *S. pombe* cells, as seen inside VR. The model was generated from a public dataset (EMPIAR: 10988)^26^ as described in ref.^27^ and displays ribosomes, nuclear pore complexes, fatty acid synthases and the nuclear envelope. Other content in the same room includes models of three different viruses and bacteriophages, with their identifying EMDB codes provided in labels (Figure 3G).

Along the lines of general chemistry, we also incorporated rooms created specifically for undergraduate students with content about the 3D shapes of atomic orbitals and about VSEPR theory, deployed during activities as explained in a separate paper^31^ (Figure 4A), and we are planning to soon add a room with a periodic table of the elements for a series of specific activities with high school and early undergraduate students (draft version in Figure 4B). Likewise, besides a standard room for structural biology, we incorporated a room specifically tailored to a MSc/early PhD-level course in integrative structural biology that covers the main techniques used to determine protein structures. In this case we used malate synthase G as a test case, as its structure was separately solved by NMR, X-ray and Cryo-EM.^28–30^ By showing these structures and some of the experimental data used to derive them, we illustrate the strengths, limitations and complementarity of the three main atomic resolution techniques (Figure 4C and 4D).

**Figure 4.**
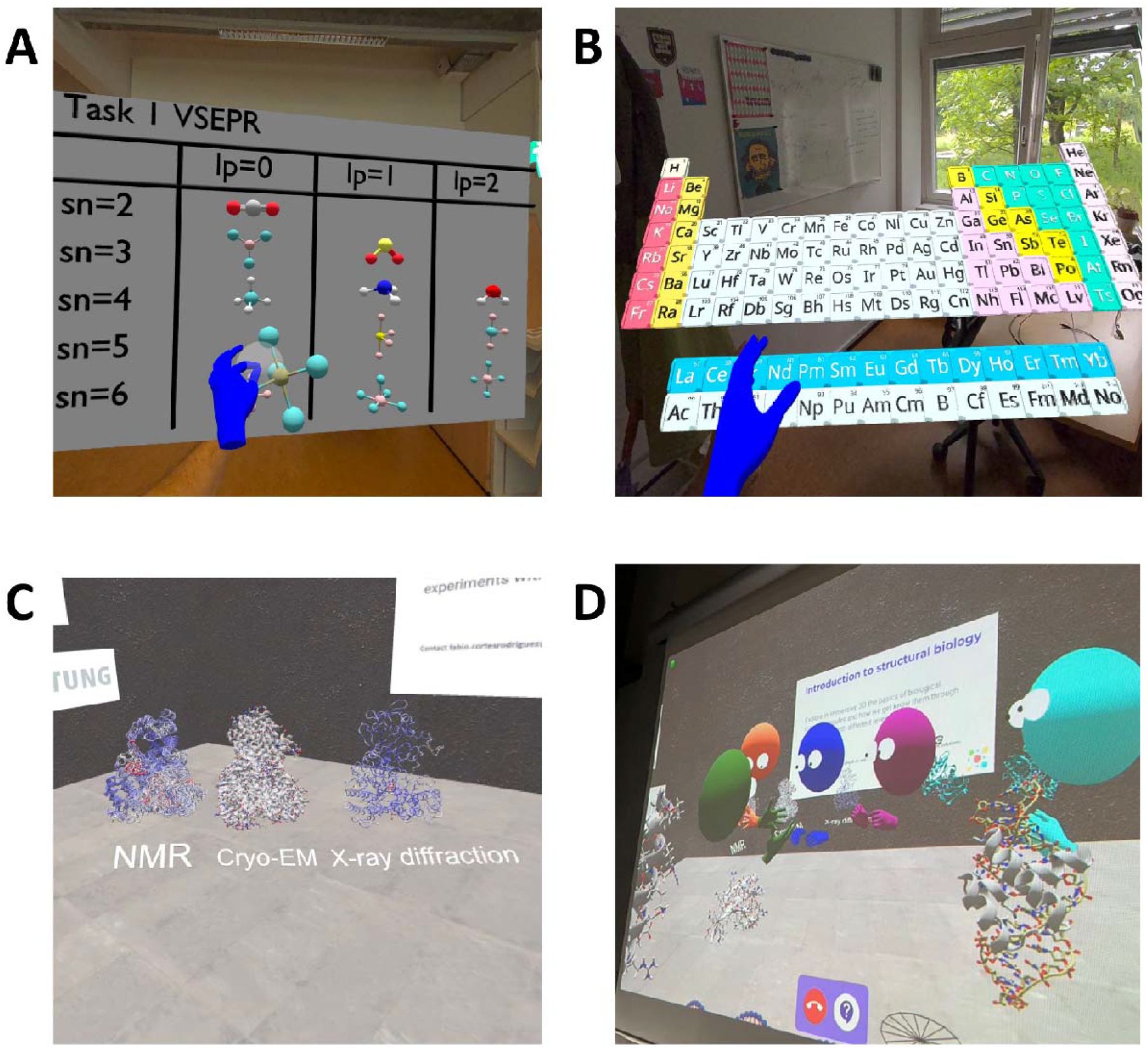
Some of the VR rooms built specifically for deployment in courses. (A) VSEPR theory for undergraduate students, utilized for lessons at Mahidol University as described in ref. ^31^. (B) A virtual periodic table of the elements, in preparation for future deployment in class. (C) Structure of Malate Synthase G as determined by NMR spectroscopy, cryo-electron microscopy and X-ray diffraction, used in a MSc / PhD level course on integrative structural biology. (D) Four students (red, blue, magenta and cyan avatars) following inside VR the explanations of an instructor (green avatar).

Besides accessing the preset content, users can create with our PDB2AR tool^23^ *ad hoc* content for their presentations starting from either raw PDB files (uploaded locally or fetched from the PDB or the AlphaFold-EBI database) or from VMD-generated wavefront objects. In this case, users must follow the procedures described in our previous work,^23^ and then copy-paste the link to the GLB file obtained via email. An example application of custom-generated content is shown in Figure 2A for which the users had to prepare virtual representations of the structures they presented as part of a series of scientific talks during a conference.

Furthermore, starting from the empty room and by leveraging content created in-house with PDB2AR and downloaded or purchased from the 3D art platform Sketchfab.com, we have held sessions designed specifically for certain science communication events and school visits where we show not only molecules but also models. For example, Figure 2D is a snapshot from a 15 min long VR session where a presenter tells an engaging story that connects physics with chemistry and biology attempting to demonstrate the continuous nature of science, with all participants inside VR.

As described throughout the paper, we have successfully applied MolecularWebXR in various formats at scientific presentations in conferences, science outreach events, and for teaching, including combinations where users wore VR headsets or employed other kinds of devices. Importantly, we have had almost 150 people trying the 15 min long VR presentation from Figure 2D (in groups of 4 to 7 people plus a presenter) and none of the participants had to quit the experience prematurely due to VR sickness or other problems. Furthermore, users who tried the experiences inside VR headsets spanned an age range from 12 to 80 years old, and all of them could seamlessly manipulate objects with their hands, even when they had absolutely no previous experience utilizing VR headsets (>80% of the participants).

## Discussion

Modern VR and AR headsets allow users to visualize and manipulate 3D models with a level of depth and realism that was unattainable just 5 years ago. Unfortunately, multiple factors hinder widespread access to the technology, leaving out a large proportion of the population as compared to solutions based on regular consumer devices. Our previous big release, MoleculARweb, tackled the need for more immersive visualization and more intuitive object control for chemistry and structural biology education through web-based but only partially WebXR-based activities and playgrounds. With a steady base of 2,500 users per month and almost 110,000 users since its launch, MoleculARweb is widely employed at schools. Now, based on similar core concepts and material but building on WebXR and with the aim of achieving more immersive sessions, we hope the new MolecularWebXR will be welcomed for education, science outreach, and scientific discussions. We note that the device we used most for development, testing and application on users, Meta’s Oculus Quest 2 with 128 GB of memory, costs under USD 300 as of June 2024, which is around that of a mid-range smartphone. In October 2023, the Meta Quest 3 was launched for under USD 600, and other companies have their own WebXR-supporting VR devices with prices starting around similar values. While these costs might still be prohibitive for many individual users, they will hopefully bring an option for VR/AR for many institutions, and will likely come further down in the future.

Profiting from the lower costs and building on web standards with cross-device/cross-OS compatibility and ease of deployment as our main values up from the early stages of the project, and by exploiting WebXR’s ability to unify all devices, we envision that MolecularWebXR can bridge the digital divide and produce more equal opportunities. It encourages a more inclusive use of VR and democratizes access to the modern possibilities that the technology offers. For example, a well-funded educational institution, company or research lab can afford multiple VR headsets to do concurrent multiuser collaboration or teaching with several students inside immersive VR; while institutions with less funding can have just one or two users wearing VR headsets to manipulate objects while other users participate from their smartphones, possibly inserted in carboard goggles to feel immersive, or simply on a computer screen or beaming on a widescreen.

We cannot emphasize enough how WebXR makes the experience highly inclusive and content so readily available. We take the chance to encourage other developers creating scientific VR/AR applications to move to this platform. All developers of VR and AR headsets, from major players like Apple, Meta and Microsoft to smaller, highly specialized companies, have included web browsers that support the standard, ensuring that developments for the web should work out of the box in all of them.^16–18^ From our side, next in the line is the release of a web app for immersive molecular simulations that will enable educational and research activities like those offered by moleculARweb’s virtual modeling kits^20^ but in multiuser, immersive VR that runs, like all of MolecularWebXR’s content, on VR and non-VR capable devices. A growing universe of tools based on WebXR has the potential to transform the way how chemistry and biology, and science in general, are taught, learned, communicated and discussed, leaving no one behind.

## Acknowledgements

This work was initiated with funds from Hasler Stiftung (Bern, Switzerland) and then executed with funds from the Swiss National Science Foundation (CRARP2_209794 and 205321_207487) to LAA. We acknowledge Prof. H. Abriel, Dr. P. Teixidor and Dr. V. Rossetti (University of Bern, Switzerland) for their help deploying VR-based scientific presentations during the closing event of NCCR TransCure. We acknowledge Dr. Ricardo Diogo Righetto (University of Basel, Switzerland) for help preparing the 3D view of *T. kivui* cell. We acknowledge funding from the Novo Nordisk Foundation (NNF22SA0076513) for support for the course on integrative structural biology at the University of Copenhagen, Denmark.

