## Supporting Information for "MolecularWebXR: Multiuser discussions about chemistry and biology in immersive and inclusive VR"

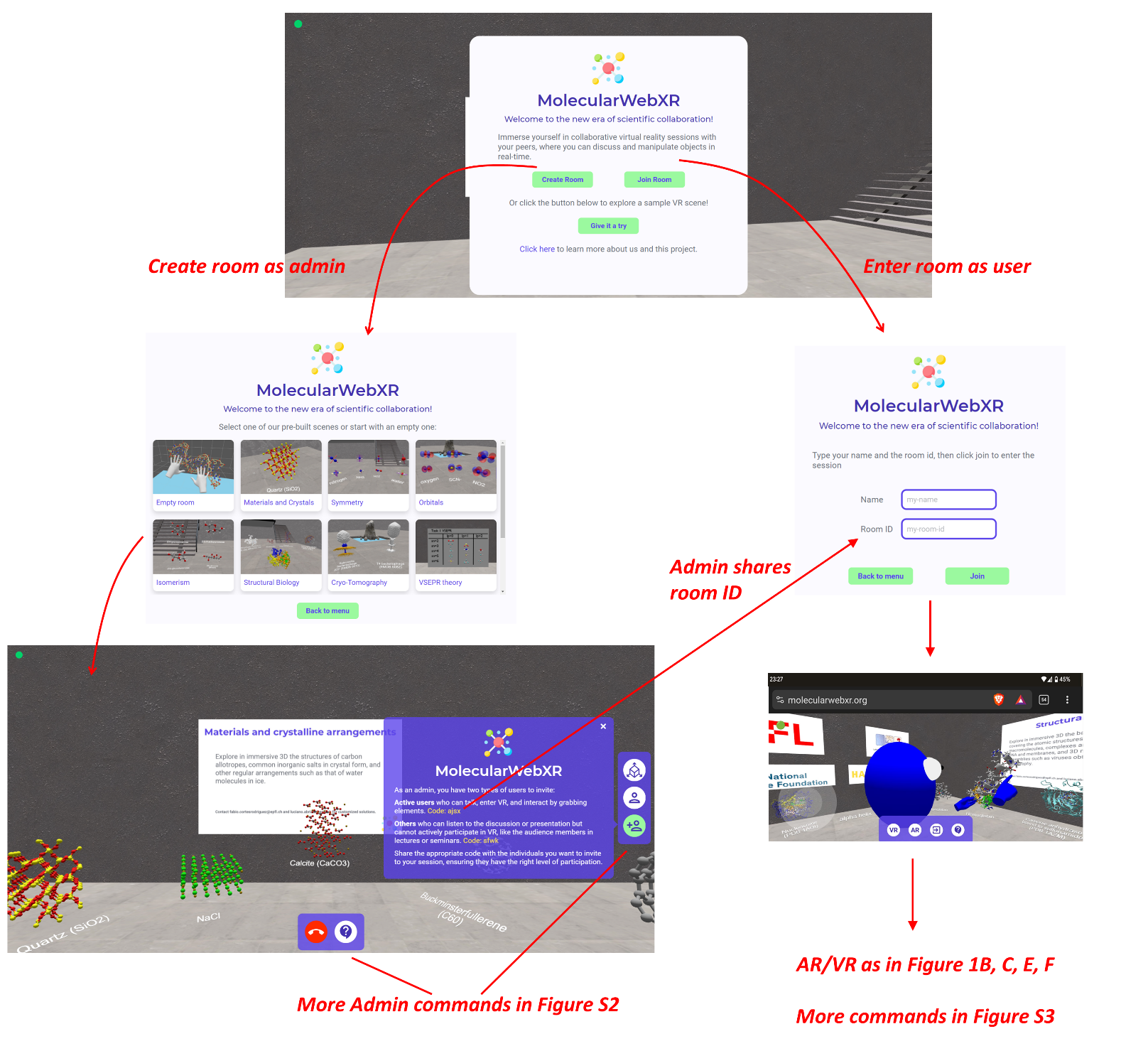

**Figure S1.** On accessing molecularwebxr.org, the user can create a new room becoming its Admin (left) or join a room created by another Admin (right). The VR and AR buttons show up only if the device and its WebXR API supports each mode; typically, modern smartphones and VR headsets allow both. Check Figure S2 for more Admin commands, figures of the paper for examples of AR and VR views, and Figure S3 for more commands available for Active users.

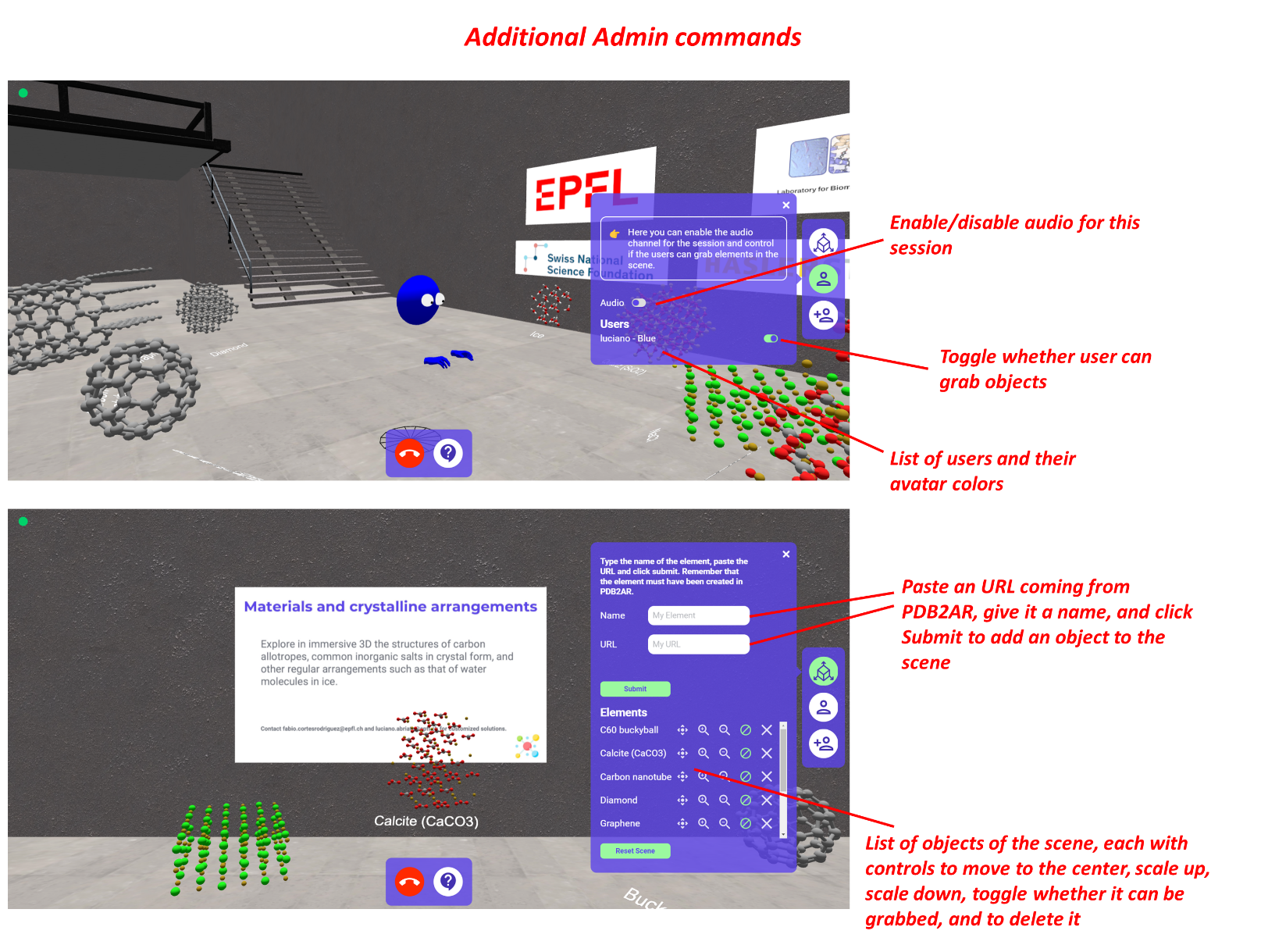

**Figure S2.** Admins can run on any device, but we advise running them on large screens such as tablets or laptops as in this view. Besides the two buttons in the middle bottom (*Hang* button in red to finish the session and *About* (?) button), Admin users have three main side menus that allow them to control virtual objects (bottom screenshot) and users (top screenshot), plus the panel showing information about the room (Figure S1, bottom left). The panel to control virtual objects allows the Admin to add objects (only accepted if created with molecularweb’s PDB2AR tool), scale them up or down, move them to the origin, delete them, or toggle whether they can be grabbed or not. The panel to control users has a switch to toggle audio functions with which users can talk with each other, and also displays a list of all active users with controls to allow them to grab or not.

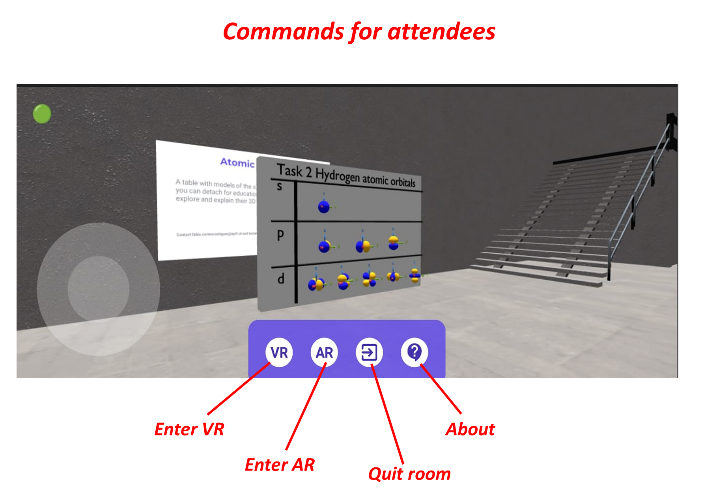

**Figure S3.** Commands available to active users who enter a room from its room ID using a AR- and VR-compatible device.
